# Modular deep learning enables automated identification of monoclonal cell lines

**DOI:** 10.1101/2020.12.28.424610

**Authors:** Brodie Fischbacher, Sarita Hedaya, Brigham J. Hartley, Zhongwei Wang, Gregory Lallos, Dillion Hutson, Matthew Zimmer, Jacob Brammer, The NYSCF Global Stem Cell Array^®^ Team, Daniel Paull

**Affiliations:** The New York Stem Cell Foundation Research Institute, New York, NY

## Abstract

Monoclonalization refers to the isolation and expansion of a single cell derived from a cultured population. This is a valuable step in cell culture so as to minimize a cell line’s technical variability downstream of cell-altering events, such as reprogramming or gene editing, as well as for processes such as monoclonal antibody development. However, traditional methods for verifying clonality do not scale well, posing a critical obstacle to studies involving large cohorts. Without automated, standardized methods for assessing clonality *post-hoc*, methods involving monoclonalization cannot be reliably upscaled without exacerbating the technical variability of cell lines. We report the design of a deep learning workflow that automatically detects colony presence and identifies clonality from cellular imaging. The workflow, termed Monoqlo, integrates multiple convolutional neural networks and, critically, leverages the chronological directionality of the cell culturing process. Our algorithm design provides a fully scalable, highly interpretable framework, capable of analyzing industrial data volumes in under an hour using commodity hardware. In the present study, we focus on monoclonalization of human induced pluripotent stem cells (HiPSCs) as a case example. Monoqlo standardizes the monoclonalization process, enabling colony selection protocols to be infinitely upscaled while minimizing technical variability.

## Introduction

The isolation and subsequent expansion of a single cell derived from a cultured population establishes monoclonality and is frequently considered an essential step in developing high-quality cell lines (Kwakkenbos *et al*., 2010). This procedure is intended to minimize or eliminate genomic and phenotypic heterogeneity in an attempt to maximize uniformity of cell lines. For instance, a newly genome-engineered cell population may comprise an admixture of cells with divergent alleles, zygosity, and epigenetic characteristics (Wang *et al*., 2017). A homogenous cell line can thus only be reestablished by ensuring all cells in the population are descendent from a single ancestral cell which was isolated downstream of any event with a high proclivity to introduce variations. This step is referred to as monoclonalization.

An example of a cell culturing process in which monoclonalization is often considered critical is in that of human induced pluripotent stem cells (iPSCs). Due to the capacity for unlimited selfrenewal and ability to differentiate via any lineage, this cell type offers immense promise for modelling disease states *in vitro*, enabling non-invasive genetic association studies, particularly as they relate to drug responses. Such efforts necessarily entail large, population-level cohorts (Vischer *et al*., 2017). Cell-line derivation throughput is therefore the paramount limiting factor in unlocking the vast promise that iPSC technology holds in relation to fields such as functional genomics and precision medicine. The iPSC reprogramming process exerts a large amount of stress on cells, resulting in a population which is highly heterogeneous with regards to variables such as residual load of viral reprogramming vector (Seki *et al*., 2012) and introduced chromosomal aberrations (Chen *et al*., 2018), eliciting the need to monoclonalize. Although Paull *et al*. (2015) described fully automated methods for iPSC production, the need for monoclonalization workflows in iPSC production remain, particularly when using viral vectors for iPSC vectors (Seki *et al*., 2012). As this step has historically incurred a critical bottleneck during automated and high-throughput derivation of iPSCs, we focus on this cell type as a case example for investigating monoclonalization methodologies.

Single-cell isolation is typically achieved via fluorescence-activated cell sorting (FACS), a form of flow cytometry (e.g. Hsieh *et al*., 2017). This process enables rapid sorting of individual cells; however there are a number means by which it can result in undesirable outcomes. Sorted cells may not survive, leaving an empty well; alternatively, faults in the sorting process may erroneously transfer more than one cell to the destination well, resulting in polyclonality. Further, for any given cell type, there may be variety of morphological or physiological changes that can occur during development that alter the quality of the cell line. In the case of stem cells, for instance, there are a number of known morphological markers which indicate loss of pluripotency (Ellis *et al*., 2009; Waisman *et al*., 2019), a common defect in newly reprogrammed iPSCs (Kyttälä *et al*., 2016; Miller *et al*., 2013). As a result of these factors, the presence, clonality and quality of cell aggregations in putatively monoclonalized wells must be validated *post-hoc*.

At present, the only method for validating monoclonality is through manual inspection of microscopic imaging performed at regular intervals to track the growth of colonies after sorting. Doing so is highly time-consuming, with technicians often spending several hours per day classifying wells according to colony presence, clonality and morphology. More critically, however, the reliance on human judgement introduces key sources of bias and technical variability, particularly when such protocols are distributed among multiple investigators and research groups. As a result of this lack of standardization, monoclonalization protocols cannot be reliably upscaled without exacerbating the technical variability of cell lines. All of these factors make monoclonalization a highly desirable target for automation, which would enable colony selection protocols to be infinitely expanded and distributed at scale while minimizing technical variability.

Deep learning, based on the use of convolutional neural networks (CNNs), has enabled enormous advances in computer vision over the past several years (Krizhevsky *et al*., 2012; LeCun *et al*., 2015) and has become an invaluable tool in automating the analysis of biomedical images of various types (Wainberg *et al*., 2018; Caicedo *et al*., 2018). These techniques have already been applied to numerous processes in stem cell research, including for the automated inference of differentiation (Kusumoto *et al*., 2018; Waisman *et al*., 2019) and prediction of function in iPSC-derived cell types (Schaub *et al*., 2019). To our knowledge, however, CNNs have never been employed in automatically identifying clonality during monoclonalization protocols for any cell type.

In domain-specific tasks, deep learning models frequently match or surpass the image-analyzing performance of human investigators (e.g. Esteva *et al*., 2017; Pereira *et al*., 2016). Dedicated neural network architectures exist for specific tasks such as image classification (Coudray *et al*., 2018) and segmentation (Havaei *et al*., 2015). Specifically, detection networks, which are trained to detect and localize each instance of a given object class in images, clearly offer a promising opportunity for the automated verification of monoclonality, which ultimately relies on the counting of individual cells. Implementations of detection networks in other scientific endeavors have previously proven highly successful (e.g. Caicedo *et al*., 2019). These typically adhere to standardized procedures for training and inference, involving annotating images with object bounding boxes for training, followed by fitting the labelled data via defined network architectures such as RCNN (Girschick, 2015) and YOLO (Redmon *et al*., 2016).

A number of key nuances inherent to monoclonalization make the task resistant to automation through standardized, widely adopted deep learning practices. For instance, confirming a monoclonal well requires the enumeration of individual starting cells. These typically occupy <0.01% of the well’s field of view and are frequently too small to be visible to human investigators without manually magnifying the image at the precise location of the cell. Grayscale imaging exacerbates this difficulty, typically exhibiting a large amount of noise. Debris particles very often appear subjectively indistinguishable from starting cells and investigators frequently rely upon information in later images, such as growth, to confirm whether a specific particle is a cell or an abiotic artefact.

Irrespective of the above, verifying clonality necessarily depends upon the interaction between images taken at different time points. For instance, enumerating individual cells in a day 0 image to validate that the sorting process was successful in isolating exactly one starting cell provides no information about the cell’s subsequent survival, expansion, or retention of desirable morphological traits. Conversely, validating that only a single colony is visible at time of inspection does not suffice to confirm monoclonality, given multiple starting cells may give rise to a single, polyclonal mass of cells which superficially resemble monoclonal colonies. In short, insofar as human investigators are able to assess, there are no cases in which a single image may contain all the information necessary to infer the clonality of a well. For this reason, it is not feasible to construct a conventional training set consisting simply of images and their corresponding semantic labels.

Here we report an algorithm design which overcomes these difficulties by leveraging the chronological directionality inherent to the cell culturing process. Our computational workflow, termed Monoqlo, integrates multiple CNNs, each having its own “modular” functionality. Monoqlo provides a highly scalable framework, capable of analyzing datasets numbering in the tens of thousands of images in under an hour using commodity hardware. Through the combination of automated stem cell culture and deep learning, this work demonstrates the first example of machine learning being applied to the identification of monoclonal cell lines from brightfield microscopy.

## Results

### Neural network modularity

We modularize the task of automatically assigning clonality into four distinct deep-learning-enabled functionalities (Fig. 1). The decision to modularize was based on empirical inferences made during preliminary investigations. Namely, consistent with the principles of transfer learning (Oquab *et al*., 2014), we initially suspected that a CNN’s feature-extracting capacity would be best optimized by consolidating all image types into a single training set. However, we found that networks trained in this manner performed poorly, often failing to distinguish between object classes. In particular, they often reported object types that could not feasibly occur in the image in question; for instance, detecting fully developed colonies in images generated immediately after seeding. This indicated that a single model would not perform well across the diversity of image magnifications and object classes employed during monoclonalization.

**Fig. 1.**
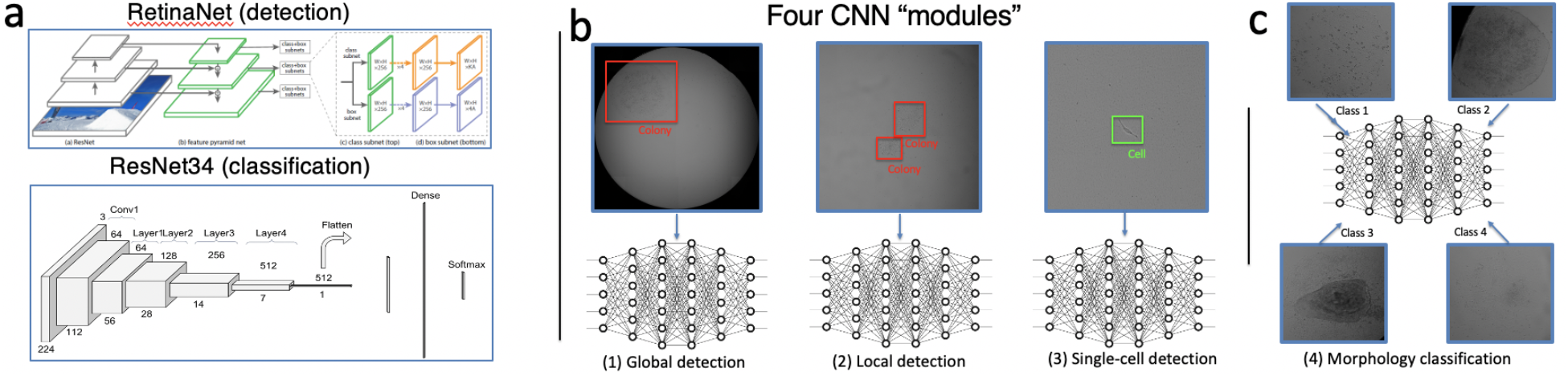
Summary of the four CNN “modules” used in Monoqlo. **a,** Simple schematic representations of the two neural network architectures used for the tasks of detection and classification. **b,** Respective functionalities of each of the 3 detection modules with representative target data and outputs. **c,** Examples of the four target morphological classes used in training the morphological classification network.

Instead, we stratify our training set based on chronological timestamps as well as magnification and crop level, and train four separate neural networks, each having its own “modular” functionality. First, we assign the term “global detection” to the task of detecting the presence or absence of colonies in a full-well image. Second, we refer to the task of detecting colonies in cropped images of various well regions at a variety of zoom magnifications as “local detection.” Third, the task of enumerating individual cells in a fully magnified, cropped image we term “single-cell detection.” We sought to achieve all three of the aforementioned tasks through the use of the Retinanet detection architecture with focal loss (Lin *et al*., 2017). Finally, in the only entirely classification-based task in this effort, we aimed to train a model to categorize images cropped around colony regions into morphological classes, here referred to as “morphological classification” (summarized in Supplementary Fig 1). Modularizing in this manner enabled us to capitalize on the temporal directionality of the cell culturing process; for instance, restricting detectable object classes to those that may realistically exist in an image based on its scan date.

### Workflow design overview

We designed a computational workflow, termed Monoqlo, that integrates each of our trained neural networks. The laboratory automation workflow that generates data for use with Monoqlo and the design of Monoqlo itself are summarized in Figs. 2 and 3, respectively. Our algorithm processes images on a per-well basis in a reversely chronological fashion. That is, for each physical well, the algorithm begins by analyzing the most recently generated scan. In our case, this is an image that has been cropped only to remove the black borders of the image, preserving the entire field of the physical well. These images are passed to the global detection model, the output of which is a coordinate vector demarcating the bounding boxes of any detected colonies.

**Fig. 2.**
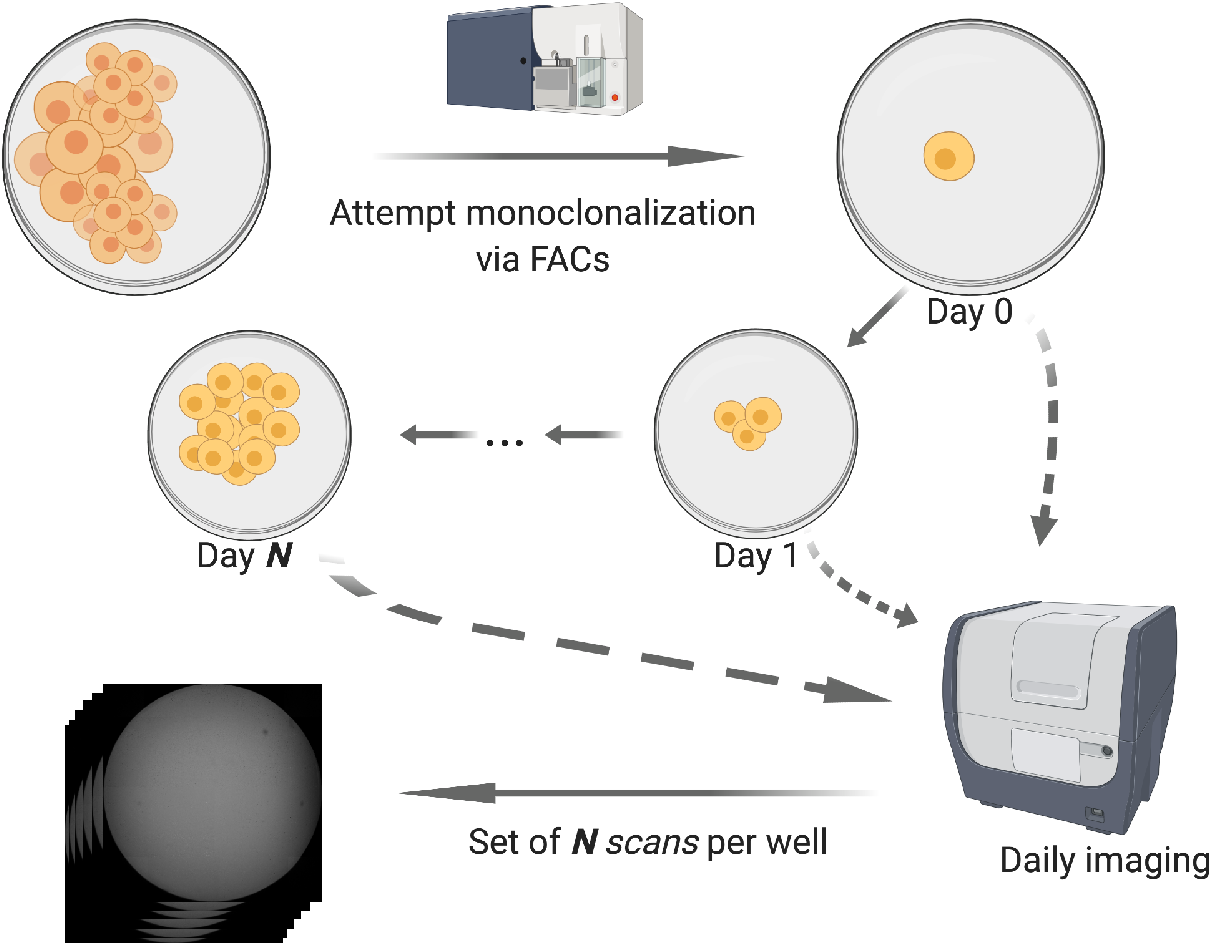
Overview of the daily automation workflow which generates data for training and real-time use with Monoqlo. Following cell deposition via FACs, cells are allowed to grow over N days, with well-level imaging occurring nightly. N represents a variable number dependent on cell growth rate and decisions on passage timing.

**Fig. 3.**
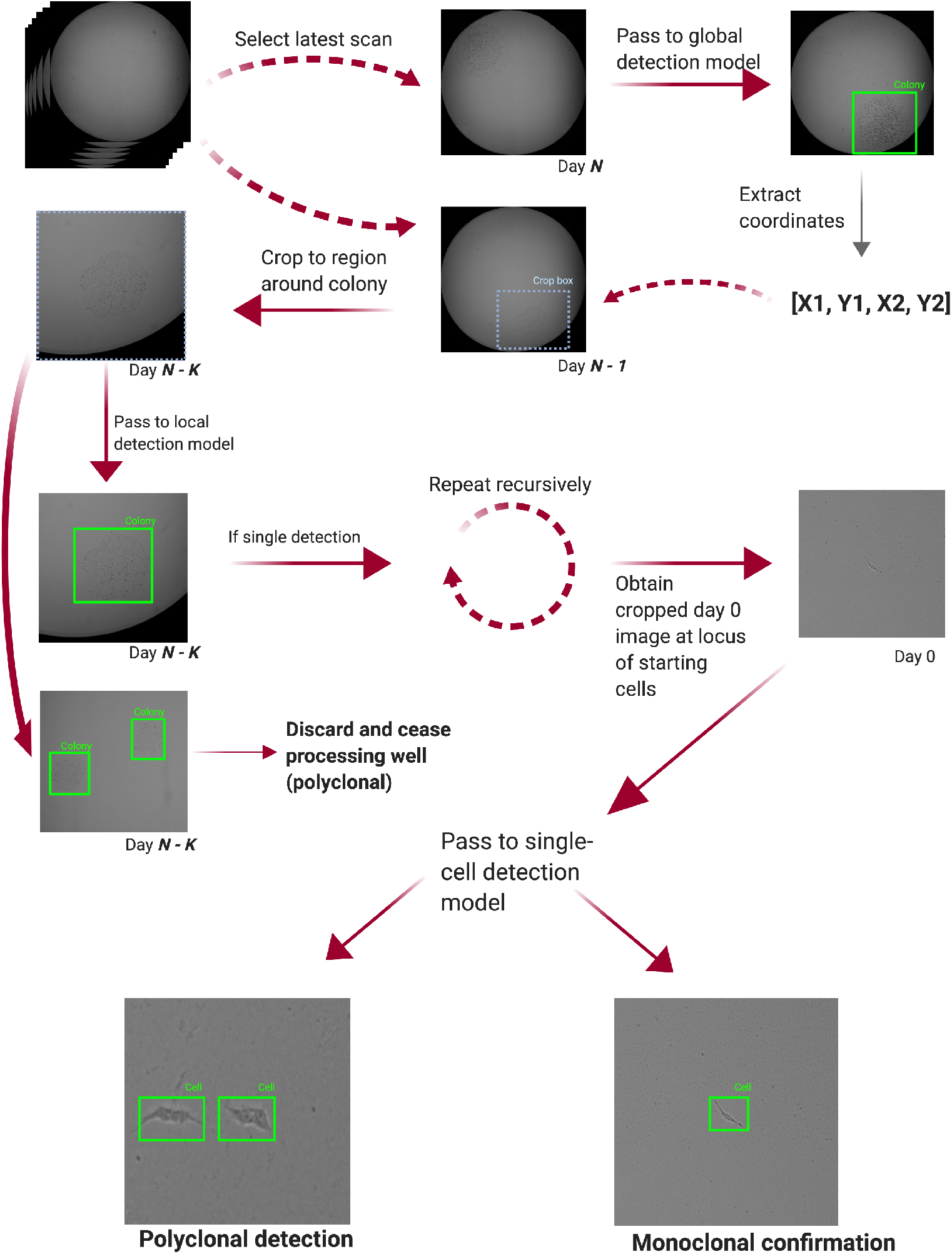
Schematic representing a broad overview of Monoqlo’s design and algorithmic logic. Arrows represent the processing order in the algorithm’s reverse-chronological analysis, beginning with the most recent scan. If a colony is detected, the region around the colony is cropped in the previous day’s scan and the image is passed to the local detection model. The process is repeated, progressively reducing the field of view being analyzed. If multiple colonies are detected in any scan, the well is declared polyclonal and no further scans are analyzed. Upon reaching the earliest “day 0” scan, the resulting image is passed to the local detection model. Based on the number of cells detected, a clonality for the well is finally declared.

Our algorithm then expands these coordinates until each dimension of the bounding box is twice that of the predicted colony, loads the next most recent image for the same well, and crops the image to the resulting region. Due to the preservation of plate orientation and physical positioning between scans, the earlier instantiation of the same colony is therefore approximately centered within the newly cropped image. The expansion of the field-of-view’s cropping coordinates beyond the size of the original bounding box simply allows for a margin of positional error. The resulting image is then passed to the local detection model, which reports the bounding box of the earlier colony, indicating its position within the original, uncropped image when summed with the cropping coordinates. The algorithm iterates this process recursively until the resultant most recent image is the earliest (“day 0”) scan, generated within hours of sorting. We found this incremental, iterative processing aspect of the workflow, as well as the expansion of the crop box dimensions, to be essential, as there are invariably small deviations from precise concentricity with each day due to non-radial growth and minor positional shifts between scans. Over periods of several days of imaging, these deviations sum to substantial offsets. As such, simply cropping and magnifying at the exact center of a late-stage colony will rarely yield a field of view in which the starting cell or cells are situated.

Aside from counting individual starting cells, polyclonality can often be inferred if two or more clearly distinct cell masses are observed, which are assumed to have originated from two or more cells from the same FACS sort. If either the global or local detection models reports a colony count of >1 at any point during the process of iterating backwards chronologically, the algorithm accordingly declares the well to be polyclonal and ceases processing any further images for that well. Alternatively, if the workflow continues to detect exactly one colony until reaching the day-zero scan, the resulting image will be magnified and cropped exactly around the ancestral cell or cells. This image can then be passed to the single-cell detection model, providing a count of the number of starting cells. On this basis, the well may then finally be declared either monoclonal or polyclonal.

### Chronological processing logic enables optimization

In our case, any given monoclonalization “run” typically comprises between 300 and 900 plate-wells and 2-6 runs are typically active at any one time. With per-well scans occurring daily for up to 30 days, the mean volume for each run at time of processing by our algorithm is therefore approximately 30,000 images. Rather than pertaining to images, however, the target labels in the case of monoclonalization correspond to individual wells. For this reason, we employ a “well knockout” approach in which detection by the workflow of any one of a number of exclusion criteria causes the algorithm to eliminate the entire well from the workflow and ignore all subsequent scans for that well. For instance, if no objects are detected in the most recent scan, then the well is reported empty at time of analysis and its antecedent characteristics are considered irrelevant. During testing, we executed Monoqlo on 8 plates of 96 wells. The mean number of empty wells per plate at time of processing was found to be 73, ranging from 41 to 92. Thus, in cases where Monoqlo is applied to, for instance, 8 plates at day 15 of the monoclonalization process, the well knockout approach alleviates the need for processing of approximately 8,760 of a total of 11,400 images (76.8%) on the basis of emptiness alone. Any well found to be polyclonal at any stage of analysis is also excluded from further processing. In the same test run, we found a mean of 11 polyclonal wells per plate, with polyclonality being declared after a mean of 5.73 images having been processed. During real time deployment, we further extend our exclusion criteria to eliminate wells which are found to exhibit the morphological markers of differentiation. Due to the enormity of the datasets that require daily analysis, our knockout approach provides a vast improvement to compute time.

### Neural networks learn to detect colonies and classify morphology

We began by evaluating learning trajectories and benchmarking the prediction performance of each CNN in its respective task. In the case of object detection networks, our initial metrics for assessment were the change in value of the loss function when tested on a held-out validation dataset representing 20% of our total image set. While training such networks, precise accuracy metrics are not automatically generated by the learning algorithms, since the model may correctly detect an object without the labeled and predicted bounding box coordinates matching exactly. As an alternative, we manually evaluated their performance by visually comparing labels and predictions in validation images with their respective bounding boxes drawn. From these comparisons, we quantified detection performance according to two metrics: 1) percentage of labelled objects which were correctly predicted and classified, and 2) number of false positives, in which the model detected an object where none was present, as a ratio to the total number of images analyzed. Results of our model validations are summarized in Fig 4. Finally, true colony width, as measured by biologists using an image scale bar, was highly predicted by Monoqlo-predicted bounding box X dimension (Pearson’s r(266) = 0.917, p < 2.2e-16) (Supplementary Fig 2).

**Fig. 4.**
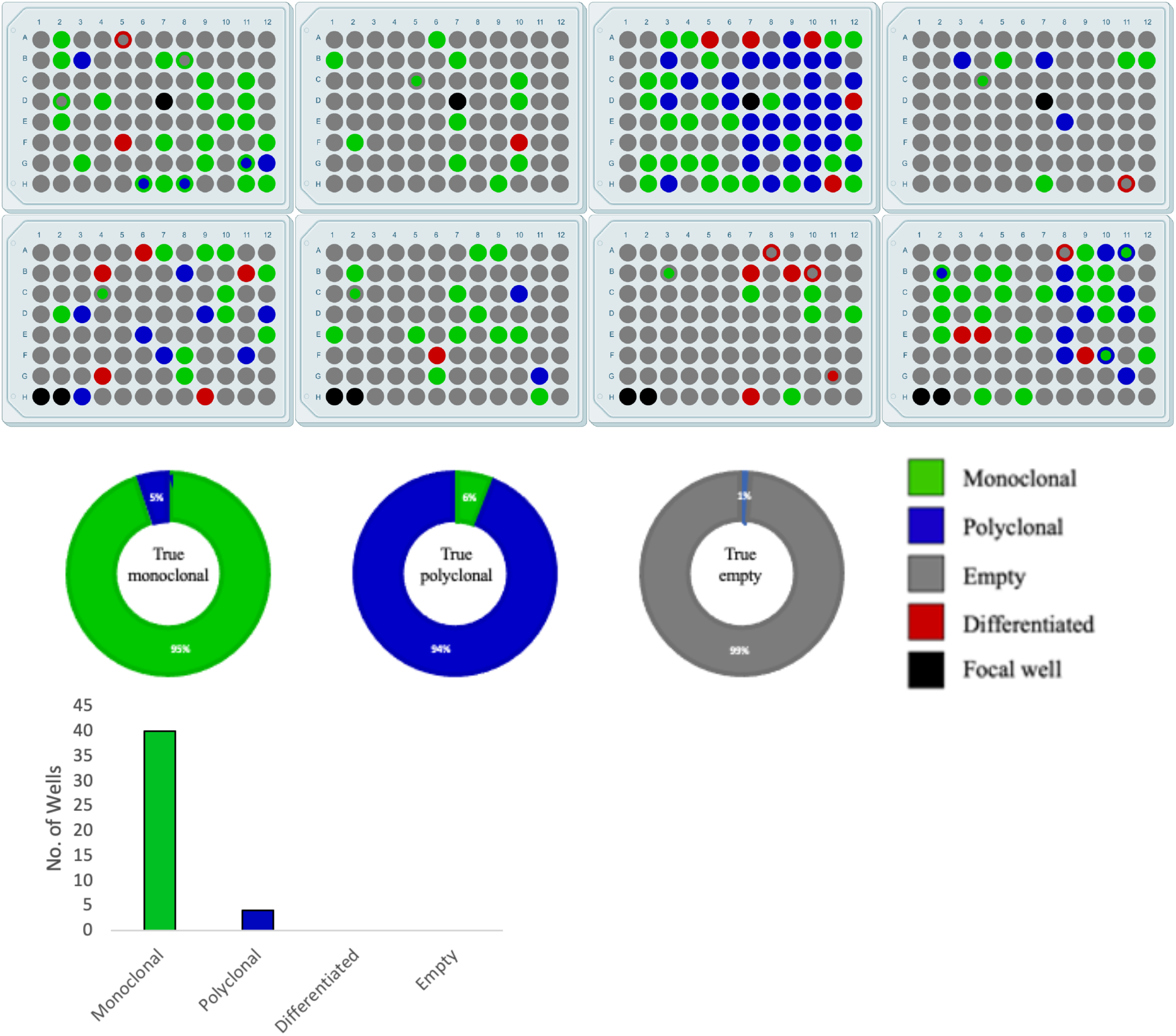
Results of Monoqlo (MQ) framework validations. **a,** Well-level clonality identification performance of the MQ framework on real-world production run data. Outer colors represent the ground-truthed clonality of the well, with color meanings indicated in legend; inner colors represent the clonality identified by MQ, with dual-color wells thus indicating MQ errors. **b,** Class-specific clonality identification performance of Monoqlo on manually curated, class-balanced test dataset. **c,** Summary of MQ clonality performance, with analyses restricted to monoclonal, morphologically healthy wells that were selected for further passaging by biologists.

### Deep-learning workflow with modularization identifies clonality

We benchmarked the efficacy of Monoqlo as a unified, modular workflow, first by testing its accuracy on a manually curated, class-balanced validation set, and subsequently by evaluating its clonality identification performance (irrespective of morphology) *post-hoc* on a raw, unfiltered dataset from real-world monoclonalization runs. Our curated test set included 100 wells from each of three classes; empty, monoclonal, and polyclonal; randomly selected from historical records of manually classified wells. The imaging date at which processing was initiated for each well was randomly generated from the range of days 8 - 18. The real-world scenario validation was performed on a monoclonalization run (DMR0001) which comprised 768 wells in total, spanning a time frame of 19 days, thus yielding a data volume of 18,240 images. Manual image review found 561 of these wells to be empty; that is, they contained no indication of living cells, irrespective of remnants of dead colonies, abiotic debris, and other artefacts. Monoqlo correctly eliminated 556 (99.1%) of these wells. The remaining 5 empty wells were reported as monoclonal, seemingly resulting in false positives on the part of the global detection model due to unidentified abiotic artefacts (Supplementary Fig 3) with a superficially similar appearance to that of a cell colony. Accordingly, Monoqlo identified 194 non-empty wells. This included 115 monoclonal declarations, of which 2 and 5 wells were found to have ground-truth classifications of “polyclonal” and “empty,” respectively; the remaining 108 wells (93.9%) being concordant with the ground truth. Finally, 61 wells were reported as being polyclonal, of which 57 (93.4%) were confirmed by ground truthing and 4 were found to be monoclonal. Results of both validations are summarized in Fig 4.

### Hand-crafted programmatic solutions improve deep learning workflows

We identified a number of circumstances in which shortcomings of our trained CNNs, which would otherwise have led to erroneous results, could be robustly corrected for using simple programmatic logic. Perhaps most prominently, we found that detection CNNs tended to often report multiple, overlapping colonies in image regions where only a single colony existed in the ground truth (Supplementary Fig 4). We found were able to partially mitigate this by adjusting the size and distribution of the anchor boxes (Jeong *et al*., 2018). However, doing so is laborious, can only be done prior to training a model, and provides only an incomplete solution. Instead, our algorithm combines any overlapping boxes and considers the resulting box as a single object. In the case of colony detection, this never results in loss of polyclonal identifications. To illustrate, consider the concept of “colony splitting” which occurs due to Monoqlo’s reversely chronological approach. Colonies which overlap one another at day ***N*** are spatially isolated at day ***N - K*** and have grown into a combined mass at day ***N + K*** where ***K*** is a variable amount of time dependent on growth rates and original separation distance (Supplementary Fig 5). Thus, overlapping object detections can safely be considered by our algorithm as a single object which, if representing multiple colonies, will later be detected as entirely isolated from one another in earlier images and thus declared polyclonal.

## Discussion

This work represents, to the best of our knowledge, the first to automate the identification of clonality using a deep learning object detection approach. We believe this has the potential to remove a critical restriction on scalability in a number of cell culturing domains. This includes the present case of iPSC derivation, where monoclonalization is considered essential for two reasons. First, in cases of viral reprogramming, there is a large amount of cell-to-cell variance in residual load of the Sendai viral vector used to deliver transcription factors to the inner cell during reprogramming (Agu *et al*., 2015). Second, the reprogramming process often leads to severe chromosomal abnormalities (Chen *et al*., 2018), presumably due to stress-induced mitotic disruptions. Both of these factors cause profound phenotypic variation, resulting in unpredictable, highly heterogeneous cell lines, eliciting the need for monoclonalization, which has historically incurred a bottleneck during iPSC production. Paull *et al* (2015) suggest that the physical monoclonalization process could exert further physiological stress on cells; however single cell cloning remains critical in a number of use cases. Given the extent to which cohort size dictates the viability of population studies, the removal of this bottleneck, as demonstrated in the present work, represents a significant step in fully unlocking the immense research potential of iPSCs.

Perhaps more significantly, in addition to initial derivation, huge efforts are being made towards optimizing CRISPR-Cas9 editing efficiency and other forms of genome engineering in iPSCs (Wang *et al*., 2017), which holds enormous potential in regard to functionally annotating gene variants (Garg *et al*., 2018), disease modelling (Rahman *et al*., 2015), and validating polymorphisms identified in genetic association studies (Warren *et al*., 2017). Due to the genomic heterogeneity the editing process introduces, newly edited populations must be monoclonalized to ensure that all cells carry the same genotypes (Wang *et al*., 2015). While we have focused on iPSCs in the present study, the same holds true for gene editing in all cell types (e.g. Chu *et al*., 2015; Smurnyy *et al*., 2014). We therefore view the genome engineering pipeline to be another critical case in which the Monoqlo framework alleviates a major bottleneck in disease research and therapeutic development.

We suspect that our algorithm could be adapted to any cell type, provided the cells are capable of being imaged and form discrete clonal masses. As an important example, antibody development is one of the most common use cases for monoclonalization (Khazaeli *et al*., 1994), due to the epitope specificity of monoclonal antibodies (Vojtěšek *et al*., 1992). Many of the most frequently used cell types in antibody development have been successfully detected in microscopy imaging with CNNs (Chen & Srinivas., 2016; Zhao *et al*., 2017). As monoclonal antibodies form the central component of many drug discovery efforts (Reichert *et al*., 2007; Nelson *et al*., 2010), the Monoqlo framework may have the potential to offer a valuable tool to the pharmaceutical industry at large.

The present study adds to previous instances of deep learning applications in iPSC process automation. In particular, there is a great deal of interest in optimizing CNNs for use with brightfield microscopy in an effort to alleviate the need for immunostaining and fluorescence microscopy imaging (e.g. Kusumoto *et al*., 2018), which comes at much larger costs to financial investment and investigation time. For instance, Christiansen *et al* (2018) successfully trained deep learning models to predict fluorescent labels from brightfield images alone. This work further demonstrates the predictive power of deep learning in various analysis tasks using simple microscopy images without the requirement of fluorescent labelling.

Waisman *et al* (2019) showed that standard CNN architectures such as Resnet50 may be trained to distinguish differentiated and undifferentiated stem cells in culture, even at early onset. Our classification CNN differs from theirs in that we stratify our training classes to a greater extent, as opposed to a binary “differentiated versus undifferentiated” approach. Doing so served to increase the robustness of our algorithm when applied in real-world cell culturing scenarios, in which there is a high degree of variability in iPSC colony morphology due to factors other than pluripotency status. Additionally, our network is trained on images cropped around distinct, singular colonies as opposed to field-of-view images containing numerous, randomly seeded cell aggregations. In this sense, our training data are more akin to that employed in Kavitha *et al* (2017), in which a vector-based CNN is used to distinguish “healthy” from “unhealthy” colonies. Further, they highlight the benefits of using segmented colony images and outline a number of key difficulties involved in doing so in an automated capacity, without which classification models using segmented colony images cannot be integrated into real-world automated production workflows. By using our classification network in conjunction with colony detection models from the wider Monoqlo framework, we automate the segmentation step, enabling fully autonomous deployment in laboratory automation scenarios.

We recognize a number of shortcomings of our approach. For instance, in cases where two or more starting cells are displayed precisely adjoining one another in the earliest available scan, the well’s clonality status must be considered ambiguous. This is because it cannot be determined whether the cells were sorted independently from the source plate or if a single cell was successfully sorted in isolation and subsequently divided. Notably, however, there is a time lag between seeding and attachment of the cell to the substrate during which the cell cannot be imaged. For this reason, the timing window of the first scan is critical. Certain other efforts have attempted to address this ambiguity through fluorescence microscopy applied to nuclear-stained images, which allows nuclear segmentation and helps to resolve the spatial distribution of individual cells. However, this does not entirely eliminate ambiguity since physically adjacent cells, whether clearly distinct or not, could certainly still have a polyclonal origin. We suggest that there are a limited number of feasible approaches to handling this ambiguity. Investigators may wish to simply assume any well containing multiple cells at time of earliest scan is polyclonal. Otherwise, we suspect that the ambiguity can only be resolved by generating images taken within minutes of seeding. Due to the time lag that occurs before cells can attach, however, we submit that optical focusing issues will be inevitable. Thus, starting cells are likely to be invisible at times, making it impossible to reliably verify monoclonality.

While our algorithm represents the state-of-the-art in automated clonality inference, our results indicate that its performance remains imperfect. As a result, the exact manner of its application, and the extent to which it can be relied upon as a fully autonomous monitoring system, will depend on the aims and priorities of the process to which it is applied. If the acceptable margin of error for misclassifying polyclonal wells as monoclonal is 0%, for example, then *post-hoc* validation by human investigators will be required. Such instances will nonetheless benefit from enormously reduced workloads for manual image review. If, alternatively, the paramount priority is that nonempty wells do not go undetected and small error rates in clonality assignment are deemed acceptable, then the critical variable will be the performance of the global detection model. In the present study, the 100% detection rate for colonies of sufficient size for passaging suggests Monoqlo’s suitability for deployment as a dependable, fully autonomous system. In both of these cases, however, the defining variable will be the performance of the deep learning models in isolation. CNNs trained by different investigators and tailored to different cell/image types could show substantially inferior or superior performance to those described in the present study.

As for future directions, we believe Monoqlo could help facilitate investigations in a number of key questions which remain to be answered with regard to the predictive potential of deep neural networks in iPSC research. A number of studies have demonstrated that deep learning approaches can sometimes discriminate between biological groups in images where a morphological phenotype was not previously known to exist; or was suspected to exist but was not visible to even a trained human investigator. For instance, Poplin *et al* (2018) showed that CNNs can predict factors such as cardiovascular disease risk, gender and smoking status from individual retinal images, none of which was previously thought to manifest morphologically in the retina. Further, in the case of iPSCs, deep neural networks have been successfully trained to predict donor identity from imaging of clinical-grade iPSC-derived retinal pigment epithelium (Schaub *et al*., 2019). With these discoveries in mind, we suggest the likely existence of thus far unidentified predictive markers in iPSC colony morphology. For instance, it may be possible to predict with better-than-random accuracy at an early stage whether a presently undifferentiated colony will spontaneously differentiate. Successfully training such a model would confer enormous benefit to iPSC derivation, given the substantial costs associated with continuing to culture cells which may ultimately become unusable. Other candidate targets for CNN classification- or regression-based prediction include Sendai virus load, future QC pass/fail status, and relative differentiation affinity for specific germ layers.

Training such models will invariably require large training volumes. The Monoqlo framework allows colonies to be algorithmically segmented and cropped from raw datasets, in addition to automatically filtering out images of empty wells which typically represent the vast majority of images. In many cases, investigators may also be able to label images in batch on the basis of the classification they assign to the most recent image of a given colony or well. Applying our classification network, which identifies differentiation, allows Monoqlo to retroactively assign labels such as “will differentiate” or “won’t differentiate” to earlier instantiations of the colony. This may mitigate the need for extensively laborious, manual reviews and labelling of unfiltered image sets, enabling partially or fully autonomous generation of large training volumes for future models. As such, our algorithm provides an invaluable tool for generating custom datasets for future investigations of the utility of deep learning in iPSC research.

In summary, we have demonstrated a framework in which deep learning algorithms with a modular design can automate the verification of monoclonality in brightfield microscopy, requiring relatively little labelling. We further expanded the functionality of our workflow to classification of colony morphology, demonstrating the potential for autonomous monitoring of monoclonal cell line development and clonal selection in automation workflows. Monoqlo represents a crucial step in enabling widespread distribution of high-throughput cell line production and editing workflows. This may eliminate a critical bottleneck in the specific case of iPSC derivation and genome editing, moving current technology closer to the goal of unrestricted upscaling and distribution of pluripotent stem cells for biomedical research applications. Finally, in contrast to depending solely on machine learning models to contend with all aspects of a given task, we view this work as a useful example to highlight the benefit of combining the now well-recognized, immense capabilities of convolutional neural networks with human-designed algorithmic solutions.

## Methods

### Monoclonalization of hiPSCs

Destination plates (PerkinElmer #6005182) were pre-coated with 17ug Geltrex™ LDEV-Free, hESC-Qualified, Reduced Growth Factor Basement Membrane Matrix (ThermoFisher #A1413302) diluted in 50uL DMEM/F12 (ThermoFisher #A1413302) for 1hr in a 37C incubator. Following incubation, 150uL of d0 media 1xDMEM/F12, 1.5x PSC Freedom Supplement, (ThermoFisher #A27336SA), 1.5xAntibiotic/Antimycotic (ThermoFisher # 15240062) and 15% CloneR™ (Stemcell Technologies #05888) was added to the 50uL of Geltrex+DMEM/F12 present in the well and incubated for 1hr in a 37C incubator. hiPSC colonies maintained on Geltrex in Freedom PSC media (FRD1) (both ThermoFisher) were dissociated with Accutase (ThermoFisher #A1110501) for 5-10 min at 37C. Accutase was quenched with Sort buffer (MACS Buffer Miltenyi, containing 10% CloneR) and the cell suspension pelleted by centrifugation at 130 RCF. Cells were stained with antibodies: SSEA4-647: 1:100; BD #560219, Tra-1-60-488: 1:100; BD #560173, CD56-V450: 1:100; BD #560360, CD13-PE: 1:100; BD #555394 before being rinsed with a second centrifugation and resuspended in Sort Buffer + Propidium Iodide (PI, 1:5000, ThermoFisher #P3566). Cells were then sorted using a FACSARIA-IIu™ Cell Sorter (BD Biosciences) into the pre-prepared destination plates using a 100μm ceramic nozzle with a sheath pressure of 23 psi. The flow cytometry gating strategy employed is summarized in Supplementary Fig 6. For samples sorted using the WOLFSorter, the Sort Buffer was supplemented with SYTOX AADvanced™ Ready Flow™ Reagent (ThermoFisher # R37173) instead of PI.

### Image acquisition and labelling

All images were sourced from repositories of historical data from the monoclonalization step employed during the iPSC production process of the NYSCF Global Stem Cell Array^®^. These images, previously used for manually verifying clonality, are generated automatically once per 24-hour period from seeding until plates are disposed of. All scans, which were generated by Nexcelom Celigo cytometers, are brightfield images at a resolution of 1 μm per pixel, providing an image dimension of 7544 x 7544 pixels after stitching from 16 individual fields. We annotated a total of 3,139 images with bounding boxes and object classes. An additional 2,224 unannotated images of empty wells were included in the training set as background-only images. During preliminary investigations, we found doing so to be pivotal in reducing the rate of false detections. All annotations were generated in Pascal VOC format using the LabelImg software (Tzutalin, 2015). We augmented our dataset by applying random flip and rotation transforms to the images (as per e.g. Perez & Wang, 2017). The morphological criteria required for categorizing each object class were designated by PhD-level biologists specializing in iPSC culture. Annotations were made by technicians of PhD-, MS- and BS-level, with all annotations being independently corroborated by an additional investigator. During initial investigations, we found that models sometimes incorrectly classified aggregations of dead cells, most of which had emanated from living colonies, as colony object detections. We corrected for this by labelling these objects as a separate class, as opposed to treating them as background. We termed this object type “overspill” (see Supplementary Fig. 7).

### Convolutional neural network architectures

For all object detection tasks in the present work, we use the RetinaNet CNN framework for one-shot detection, first introduced in Lin *et al*. (2017). The defining advancement proposed in this work was the use of the novel focal loss function, which adjusts the per-sample cross entropy loss to more heavily penalize misclassifying difficult examples than easy examples, thereby resolving the issues imposed upon object detection tasks by class imbalance. Our final detection CNN architecture consists of a ResNet50 convolutional backbone (He *et al*., 2016) with dynamic input dimensions, which performs feature extraction and passes the learned representations to a feature pyramid network (FPN) (Lin *et al*., 2017). The RetinaNet50 architecture consists of 50 convolutional blocks, each consisting of a 2D convolutional layer with rectified linear unit (ReLU) activation (Dahl *et al*., 2013) and batch normalization (Ioffe & Szegedy, 2015), and uses residual (skip) connections between several convolutional blocks. The overall network incorporates the use of anchor boxes which represent predefined candidate object locations, as described in Ren *et al*. (2015), enabling the object detection task to be trained in end-to-end fashion. The outputs from the FPN are passed to two submodels, one of which performs regression to refine the localization of the object bounding boxes, the other performing object classification. Finally, the network’s output layer produces a filtered vector denoting the posterior probability of each anchor box containing an object, including the object’s class being indicated as a one-hot vector, and the refined pixel coordinates of the predicted bounding boxes. Overall, our detection networks each have >3.6 x 10^7^ parameters. For morphological classification, we use the ResNet34 architecture. For full details on the neural network architectures, see Supplementary Materials 1.

### Training of deep learning models

RetinaNet detection models were trained using a Keras RetinaNet implementation (https://github.com/fizyr/keras-retinanet) with the ResNet50 convolutional backbone having weights pretrained on the ImageNet dataset. Preprocessing involved subtracting ImageNet means from images and normalizing pixel intensity values to the range between 0 and 1. We also implement a hand-crafted algorithm for cropping the thick black borders around the well from the image which removes the outermost line on each edge of the image and repeats until the maximum, raw pixel intensity value for the given line exceeds 70. Each CNN model was trained for 60 epochs, with weights being saved after each epoch, allowing the checkpoint with the smallest validation loss to be selected as the final model for use in the Monoqlo framework. In each case, we use the Adam optimizer (Kingma & Ba, 2014), selecting an initial learning rate of 10^−3^, reducing the learning rate by a factor of 10 every 10 epochs after epoch 40.

## Supporting information

Details on CNN architectures

## Acknowledgements

We would like to thank the members of the NYSCF leadership team, specifically R. Monsma, S. Noggle, R. Aiyar, C. Anzel, L. Schwarzbach, J. Wallerstein and S. Solomon for their support throughout this work. The authors would like to thank L. Mehran and M. Berliss for their guidance on reporting of biological research protocols. The authors would like to acknowledge H. Gaiser for his excellent implementation of the RetinaNet framework. We thank C. Richardson for his hugely helpful guidance on the release of Monoqlo.

## Author Contributions

B.F., D.P. and Z.W. conceptualized the Monoqlo framework, including the use of reverse-chronological analysis for the assessment of clonality. B.F. trained and validated RetinaNet detection models and wrote the Python software for the execution and automated deployment of Monoqlo including data-handling logic, image processing and integration of deep learning models. B.F. conceptualized the use of classification networks in automatically assigning morphological classifications to most recent colony images. B.F., S.H., B.H., D.P. and J.B. conceptualized the labelling system for classifications of colony morphology. S.H. labelled training data and trained and validated all morphology classification models. G.L. and D.P. developed NYSCF’s iPSC monoclonalization laboratory-automation and colony selection protocols. B.F., B.H., J.B., D.P. and NYSCF Global Stem Cell Array^®^ Team performed image annotations for training RetinaNet models. D.H., B.H., M.Z., J.B. and NYSCF Global Stem Cell Array^®^ Team performed physical monoclonalizations, validation of the Monoqlo framework and subsequent cell culture and imaging using robotic systems.

## Data Availability

All images from DMR0001, the full monoclonalization run used in validation of Monoqlo during this study, are available at https://storage.googleapis.com/dmr0001_imaging_data/DMR0001.zip. All images will be de-identified with regard to cell line donor and NYSCF internal logistical identifiers such as run number or project code. While NYSCF maintains copyright to the collection, these images are freely available for non-commercial, scientific research purposes.

## Code Availability

Code will be made fully available upon publication for all non-commercial purposes.

## Competing Interests

Patent pending. The authors declare no other competing financial interests.

