## Supplemental figures for "Modular deep learning enables automated identification of monoclonal cell lines"

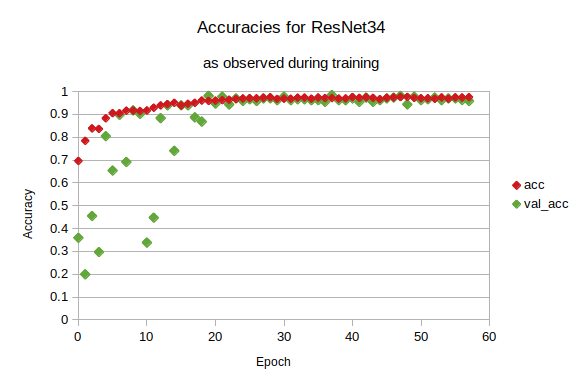

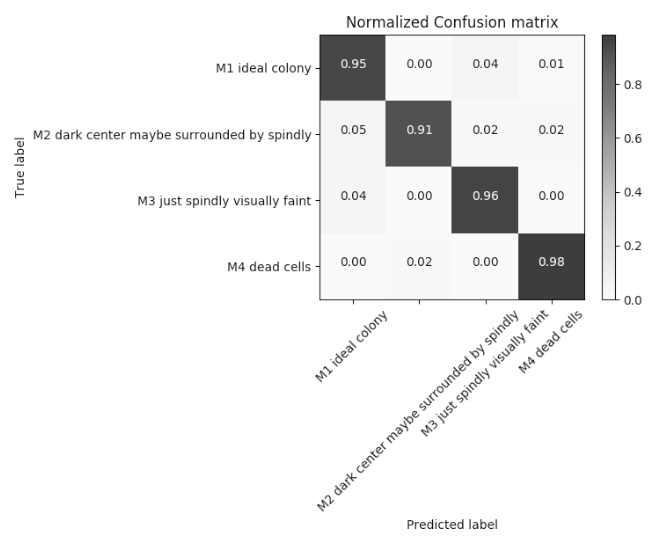

**Supplementary Fig 1 | Summary of classification model training and performance. a**, Training and validation accuracy trajectories of the classification CNN, plotted against epoch. **b,** Confusion matrix of fully trained classification CNN when tested on held-out validation set

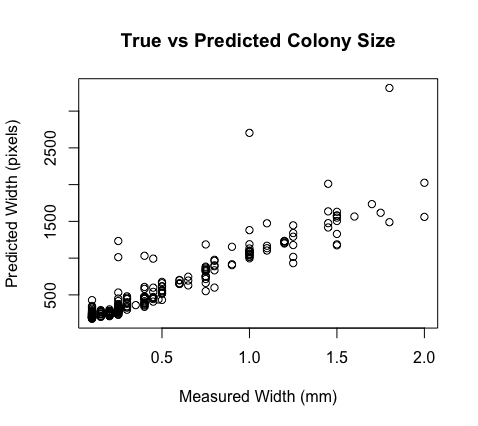

**Supplementary Fig 2**. Relationship between width of colony bounding box predicted by Monoqlo’s global detection model and the true width measured by biologists with a scale bar image overlay

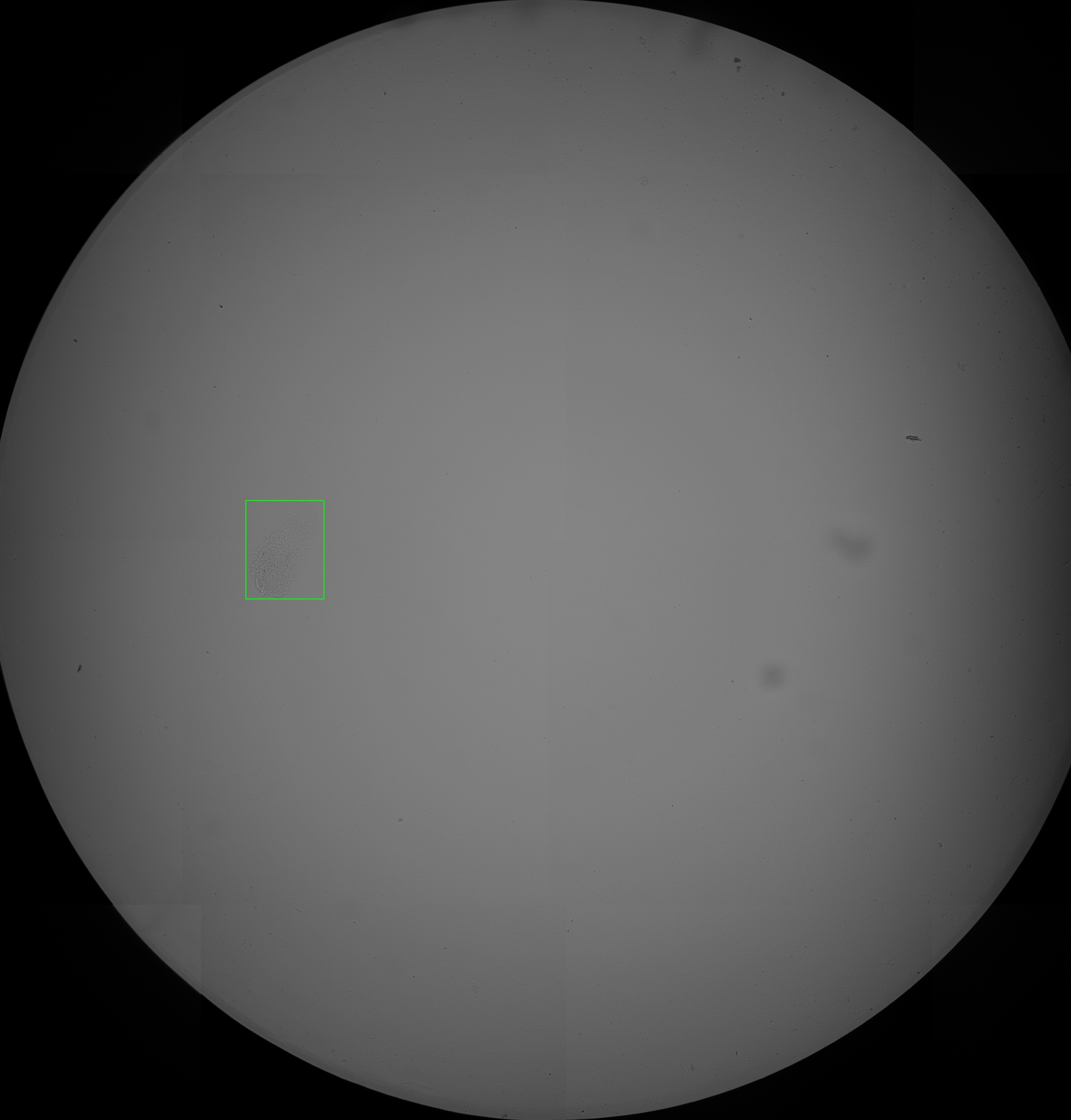

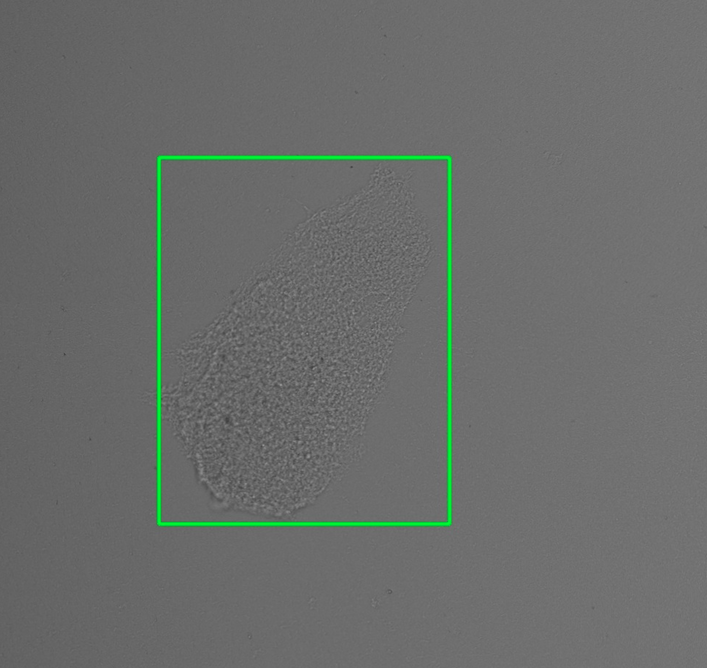

**b)**

**Supplementary Fig 3**. Example of abiotic artifacts causing false colony detections by Monoqlo’s global detection model. a) and b) represent the same image report by Monoqlo, full view and zoomed, respectively.

**a)**

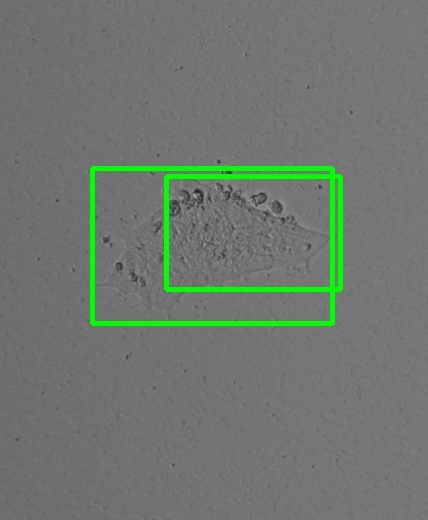

**Supplementary Fig 4**. Example of overlapping reports of colonies by Monoqlo’s local detection model where only a single colony exists after ground-truthing

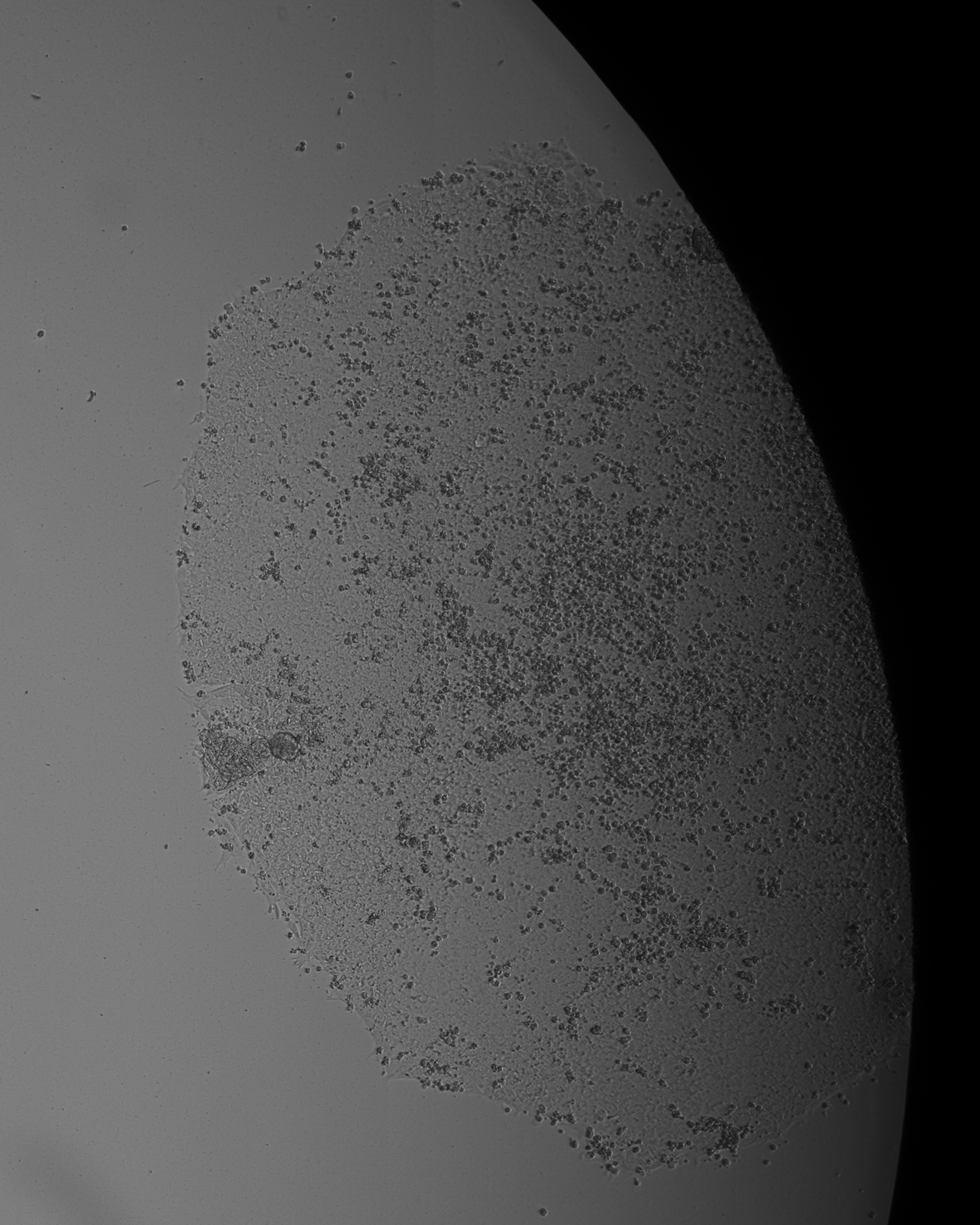

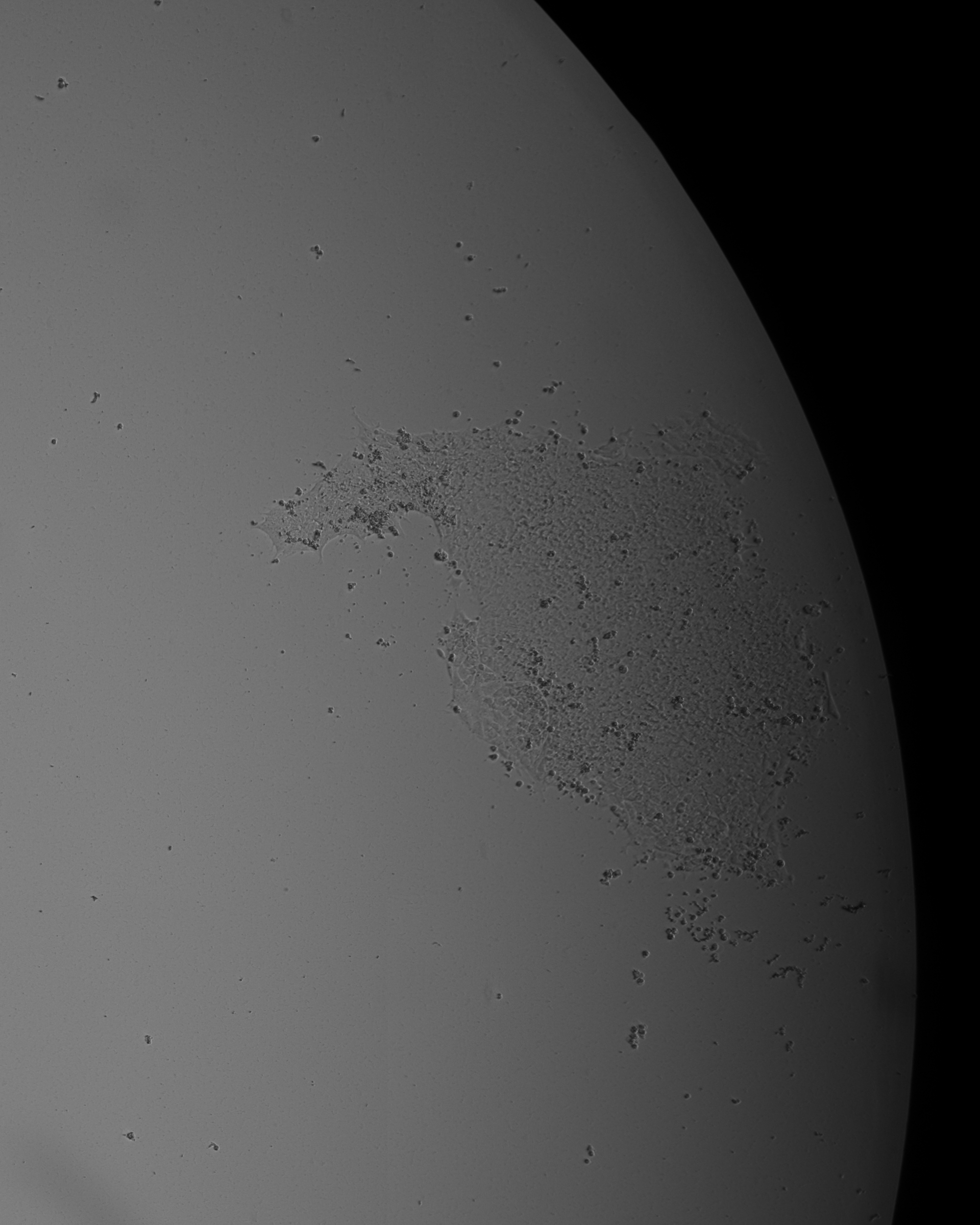

Day 8

Day 11

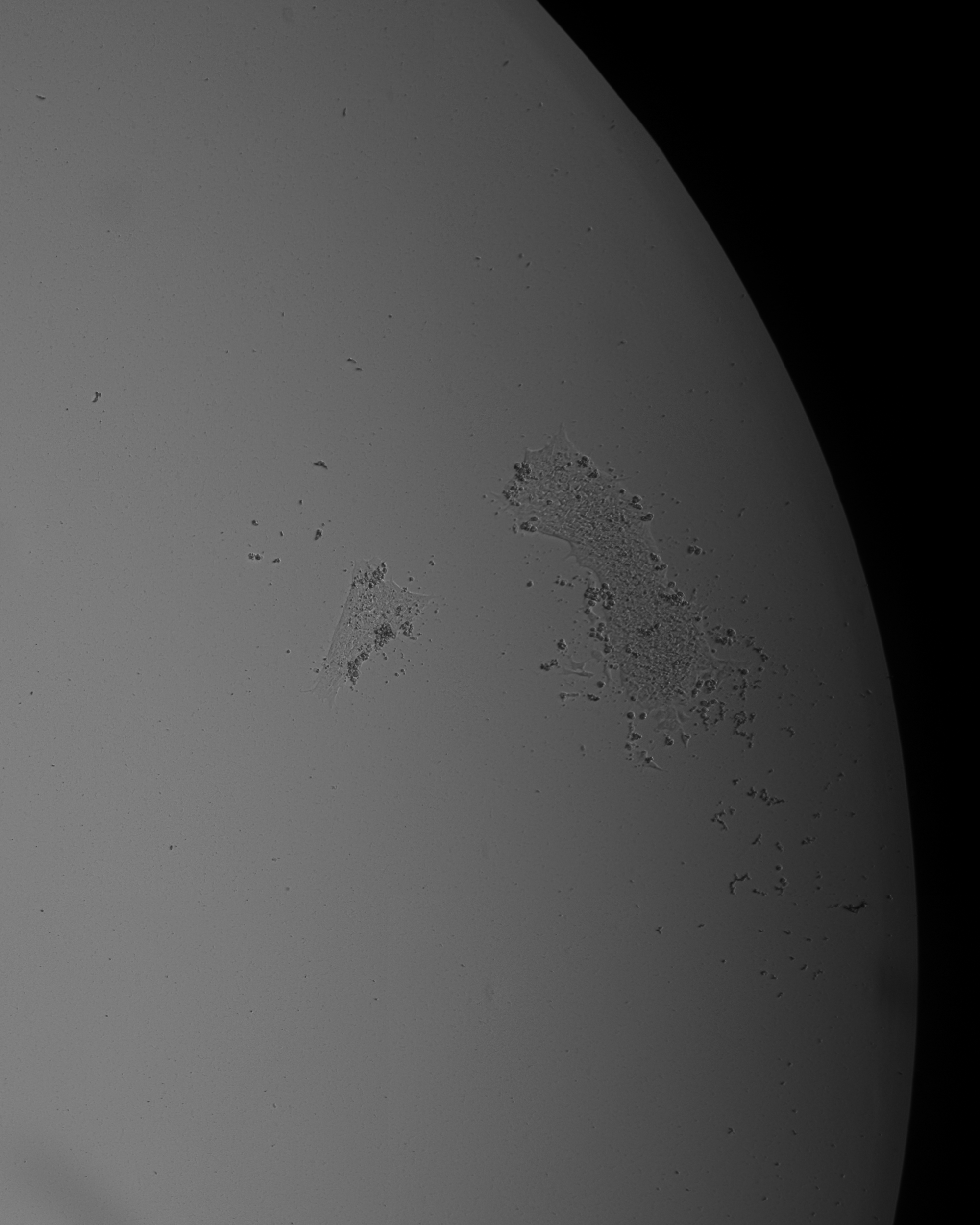

**Supplementary Fig 5**. Illustration of the concept of “colony splitting”, where an apparent single colony is revealed, during reverse-chronological analysis, to have originated in multiple colonies which ultimately merged

Day 6

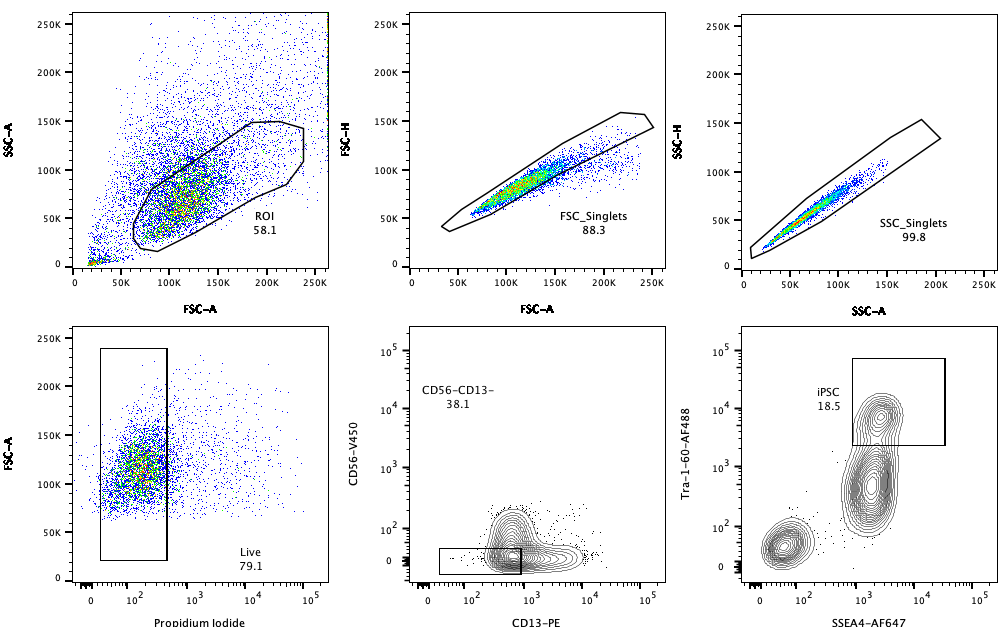

**Supplementary Fig 6**. Representative gating strategy employed during FACS-sort monoclonalization of iPSCs

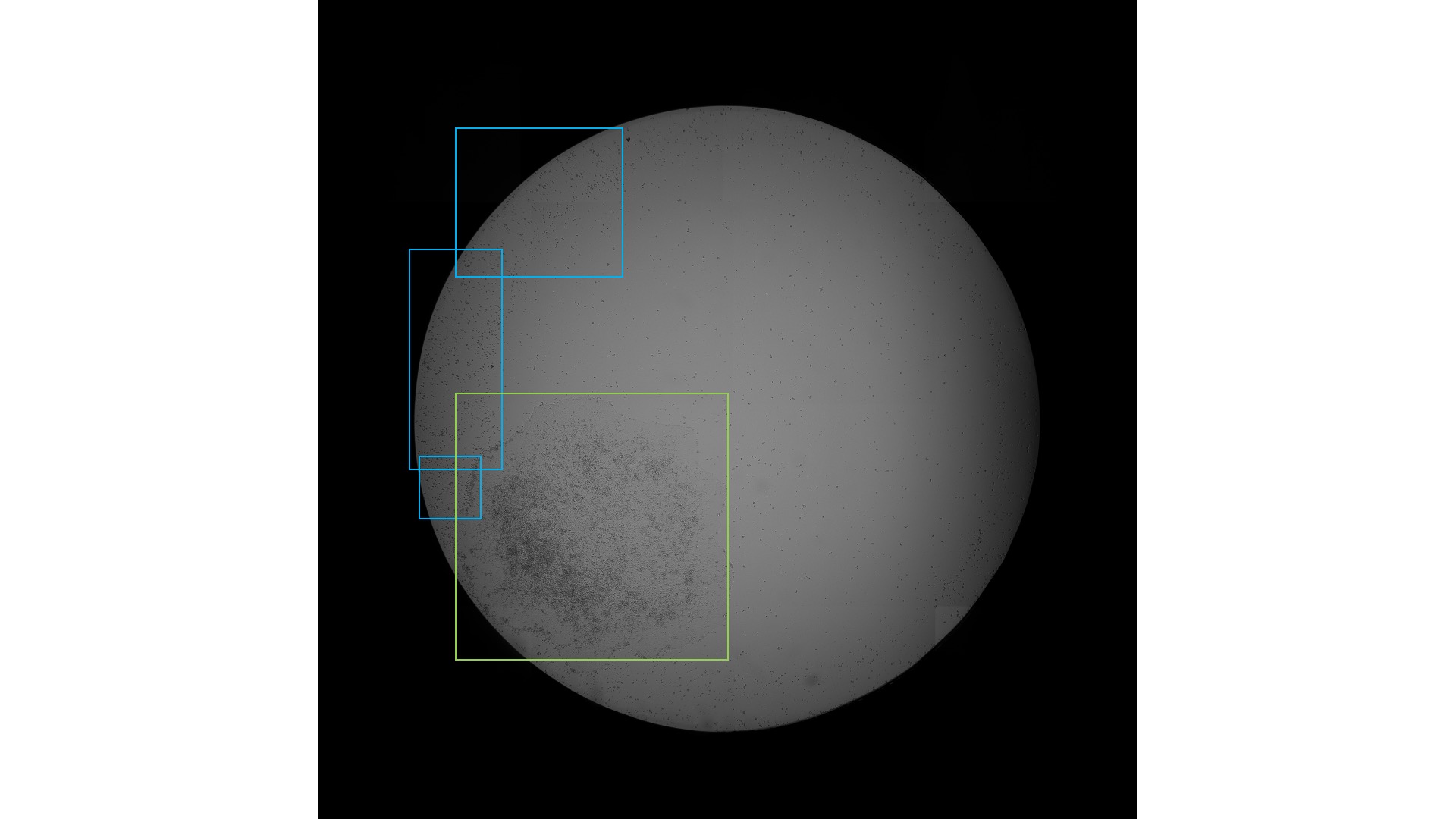

**Supplementary Fig 7**. Labelling example in which an additional object class, “overspill” (indicated by blue bounding boxes,) is annotated to improve model performance and mitigate erroneous detections of the “colony” (green bounding box) object class
