## Supplementary figures and images for "Modular deep learning enables automated identification of monoclonal cell lines"

### global_detection_model_architecture.png

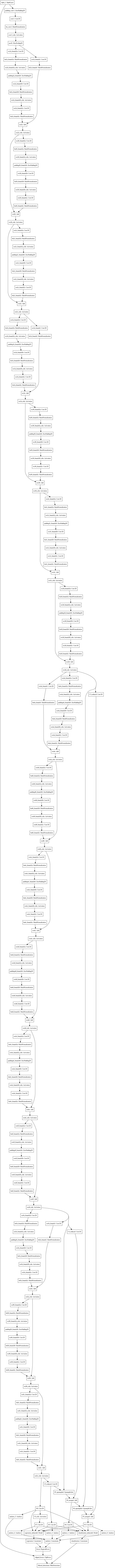

### morphology_classification_model_architecture.png

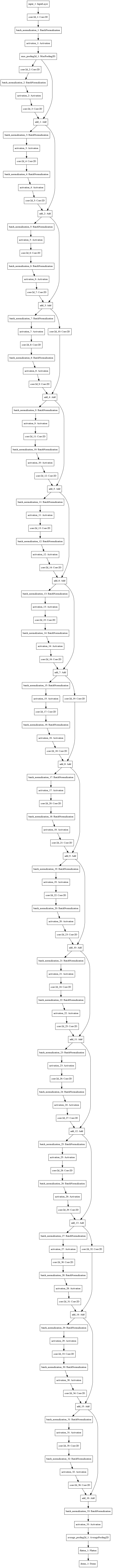

### singlecell_detection_model_architecture.png

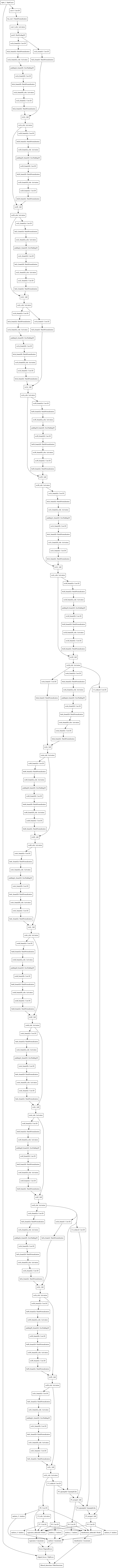
